# Longer chronic cannabis use in humans is associated with impaired implicit motor learning and supranormal resting state cortical activity

**DOI:** 10.1101/2025.04.16.647328

**Authors:** Shikha Prashad, Andrew Y. Paek, Lisa R. Fournier

## Abstract

Chronic cannabis use is associated with cognitive impairment, but its impact on implicit motor learning is unclear. Implicit learning of movement sequences (i.e., their specific ordinal and temporal structure) is vital for performing complex motor behavior and lays the foundation for performing daily activities and interacting socially. We collected data from 30 individuals who used cannabis regularly and 32 individuals who did not use cannabis. We utilized the serial reaction time task to assess implicit motor sequence learning and the Corsi block-tapping test to assess visuospatial short-term and working memory. We also recorded resting state electroencephalography (EEG). While implicit motor learning was evident at the group level, longer cannabis use was associated with a smaller index of motor learning and increased activity in beta and gamma EEG frequencies during resting state. The cannabis group also had a significantly shorter Corsi span (in both forward and backward conditions). These findings indicate that longer chronic cannabis use is associated with impaired implicit motor learning that may be a function of increased resting state neural oscillatory activity, resulting in increased cortical noise, and reduced visuospatial short-term and working memory. These findings suggest that chronic cannabis use may disrupt corticostriatal pathways that underlie implicit motor sequence learning, indicating a more extensive effect of cannabis on the motor system.

## Introduction

Recreational cannabis is legal in 23 states in the United States and is widely used around the world [1]. Acute and chronic cannabis use is associated with impaired cognition, including attention and working memory [2–6]. These cognitive processes underlie motor learning [7–9], which involves combining actions into specific temporal and ordinal structures that form complex motor behaviors critical for daily living. These behaviors include communicating with others, operating equipment and software (e.g., driving a car, using a computer), and engaging in activities that improve our health (e.g., physical activity, sports). Thus, motor learning is critical for human behavior throughout the lifespan.

Acute and chronic cannabis use may influence motor learning through movement pathways in the corticostriatal network that are modulated by dopamine. The D1 subtype of dopamine receptors modulates the direct pathway that facilitates movement, while the D2 subtype modulates the indirect pathway that inhibits movement. A careful balance between these pathways is required to perform the desired movement and suppress unnecessary or redundant movements. In addition to these movement pathways, dopamine also regulates the reward circuit from the ventral tegmental area to the nucleus accumbens and prefrontal cortex [10]. Dopamine pathways in the prefrontal cortex integrate rewarded motor behavior that leads to goal-directed actions [11].

Alterations in these pathways are associated with substance use, including cannabis, and are foundational to the development of substance use disorders [12]. For example, chronic cannabis use results in decreased dopamine synthesis in the striatum [13,14], which may impact movement pathways that connect through the striatum via dopamine projections. In addition, cannabis predominantly affects D1 receptors [15] and influences the binding of stimuli-response pairs [16] that are required for integrating task-related stimuli and appropriate motor responses. Furthermore, endogenous cannabinoids are naturally present throughout the brain and bind with cannabinoid type-1 (CB1) and cannabinoid type-2 (CB2) receptors. Animal studies demonstrate that CB1 receptors are most densely present in the basal ganglia, cerebellum, and hippocampus [17], can alter dopamine, GABA, and glutamate availability [18], and can regulate motor activity [19].

When delta-9-tetrahydrocannabinol (THC), the primary psychoactive ingredient in cannabis, interacts with these receptors, it can inhibit synaptic transmission [20]. Chronic cannabis use was found to reduce CB1 receptor availability in rats [21–24] and humans [25–27]. In rats, acute cannabis activated CB1 receptors in the cerebellum and resulted in impaired motor coordination [28,29]. These findings provide converging evidence that cannabis influences the motor system.

In addition, the effect of cannabis on dopamine and cannabinoid receptors may also alter resting cortical activity [30,31], which can then impact cognition [32]. Neural oscillations present during resting state, i.e., when an individual is awake, but not engaged in a task [33,34] have both spatial and temporal synchrony [35,36]. This neural synchrony can be measured via electroencephalography (EEG), in which neural signals in different frequencies are associated with task-specific cognitive processes. For example, alpha frequencies (8-12 Hz) are synchronized (i.e., they are present with greater power) during resting state, while beta (13-30 Hz) and gamma (30 Hz and above) are synchronized when engaged in a task. Beta frequencies in particular are associated with motor processing [37,38] in the central and parietal areas of the cortex [39,40]. Typically, resting state measured via EEG consists of greater activity in the lower frequencies (i.e., delta, theta, and alpha) and reduced activity in the higher frequencies (i.e., beta and gamma) due to the absence of engagement in a task [35]. Changes in these neural patterns may indicate disruptions in networks underlying cognition [32].

Prior studies have found that alterations in resting state EEG are associated with chronic cannabis use. For example, Prashad and colleagues reported that individuals who chronically used cannabis exhibited increased beta activity and decreased delta (0-4 Hz) and theta (4-7 Hz) activity during resting state compared to controls who did not use cannabis [41]. This altered resting state cortical activity suggests a failure to inhibit excessive cortical noise, which may disrupt cognitive and motor processes. In addition, Böcker and colleagues found greater resting state theta associated with impaired working memory after acute cannabis intoxication. They further demonstrated a dose-dependent effect of THC on beta frequencies, such that individuals who consumed larger doses of THC exhibited greater resting state beta power [42]. These findings indicate that cannabis can disrupt resting state neural oscillations that may be associated with cognitive impairment.

Building on evidence that chronic cannabis use may disrupt pathways involved in motor control and cognition, it is critical to examine whether these effects extend to implicit motor learning, which enables the acquisition and automation of activities of daily living. When these processes are compromised, individuals may struggle to learn new motor skills or adapt to novel situations, limiting their ability to interact effectively with tools, devices, and environments, and ultimately reducing independence. Elucidating the connection between cannabis use and implicit motor learning may inform public health strategies and policy. Moreover, since chronic cannabis is associated with alterations in brain activity [41,43,44], identifying neural patterns associated with motor learning impairment may yield critical insights into the mechanisms underlying cannabis-related deficits.

In this study, we examined the effect of chronic cannabis use on implicit motor sequence learning and resting state cortical activity. Implicit motor learning was assessed using the serial reaction time (SRT) task and cortical activity was measured during resting state with EEG. We also measured visuospatial short-term and working memory using the Corsi block-tapping test. We predicted that individuals with chronic cannabis use would exhibit increased resting state cortical activity (i.e., higher activity in the beta and gamma bands and less activity in the delta, theta, and alpha bands) compared to individuals who do not use cannabis. Moreover, we expected that increased years of chronic cannabis use would be associated with increased resting state cortical activity as well as larger declines in implicit motor learning. This prediction is based on prior research that indicates that longer duration of cannabis use is associated with greater neuropsychological decline [45]. Finally, consistent with the current literature [2,4], we expected that chronic cannabis use would be associated with shorter forward and backward Corsi spans (i.e., number of items in memory), indicating reduced visuospatial short-term and working memory, respectively.

## Materials and Methods

### Participants

We recruited 72 participants from the Washington State University (Pullman, WA, USA) community between March 11, 2022 and April 29, 2023. Of these, five participants did not meet the inclusion criteria and five participants did not complete all sessions of the study. Of the remaining 62 participants, 30 participants used cannabis at least four times a week for at least one year (cannabis group; mean age = 20.5 ± 1.7 years; 20 female participants) and 32 participants used cannabis 25 times or fewer in their lifetime (control group; mean age = 20.7 ± 3.0 years; 23 female participants). We used GPower [46] to estimate the sample size for each group based on *a priori* analysis using effect sizes reported in prior research that examined similar EEG outcomes in individuals who use cannabis [41]. We determined that a sample size of 30 participants in each group would be sufficient to detect an effect size of 0.6 with a power of 0.8 and alpha set at 0.05.

Participant demographic information is displayed in Table 1. Of the 32 participants in the control group, 16 reported never using cannabis, and the remaining 16 had used cannabis an average of 4.7 ± 7.4 (range: 1-25) times in their lifetime. All participants were right-handed, proficient in English, did not have a history of neurological diagnoses, and provided their written informed consent. All procedures were approved by the Institutional Review Board at Washington State University.

**Table 1.** Participant demographics (mean ± SD)

| | Control Group | Cannabis Group | $p$-value |
| --- | --- | --- | --- |
| N | 32 | 30 | - |
| Age (years) | 20.7 ± 3.0 | 20.5 ± 1.7 | 0.77 |
| Gender (M/F) | 9/23 | 10/20 | 0.66 |
| Years of education | 14.4 ± 1.1 | 14.1 ± 0.37 | 0.14 |
| Cannabis use days in prior 30 days | 0.0 ± 0.0 | 25.4 ± 4.2 | < 0.001* |
| Years of cannabis use | 0.0 ± 0.0 | 3.3 ± 1.6 | < 0.001* |
| Age at cannabis use onset (years) | - | 16.4 ± 2.1 | - |
| Lifetime cannabis use | 4.7 ± 7.4 | 2518.8 ± 2758.3 | < 0.001* |
| Grams of cannabis used per day | - | 2.3 ± 4.8 | - |
| MPS | 1.0 ± 1.7 | 4.6 ± 3.5 | < 0.001* |
| CUDIT-R | 1.1 ± 1.6 | 14.1 ± 4.3 | < 0.001* |
| AUDIT | 3.8 ± 4.8 | 7.0 ± 4.9 | 0.010* |
| FTND | 0.0 ± 0.0 | 0.0 ± 0.0 | n/a |
| DASS-21 | 10.5 ± 8.1 | 11.1 ± 9.3 | 0.81 |
| DASS-21 Depression score | 3.8 ± 3.2 | 3.7 ± 3.8 | 0.99 |
| DASS-21 Anxiety score | 2.7 ± 3.1 | 3.7 ± 3.4 | 0.21 |
| DASS-21 Stress score | 4.1 ± 3.3 | 3.6 ± 3.7 | 0.58 |
| Cognitive Failures Questionnaire | 59.4 ± 18.6 | 50.0 ± 23.0 | 0.081 |
| MCQ-SF (Session 2) | 18.7 ± 8.0 | 38.1 ± 13.3 | < 0.001* |
| MCQ-SF (Session 3) | 16.3 ± 6.8 | 34.8 ± 11.5 | < 0.001* |
| MoCA | 26.9 ± 2.2 | 27.2 ± 1.5 | 0.55 |

Abbreviations: M/F, male/female; MPS, Marijuana Problem Scale; CUDIT-R, Cannabis Use Disorder Identification Test-Revised; AUDIT, Alcohol Use Disorder Identification Test; FTND, Fagerstrom Test for Nicotine Dependence; DASS-21, Depression, Anxiety, and Stress Scale – 21 Items; MCQ-SF, Marijuana Craving Questionnaire-Short Form (not assessed in Session 1); MoCA, Montreal Cognitive Assessment.

Participants were asked to use cannabis as was typical for them (i.e., to not abstain from cannabis use prior to any of the sessions) in order to assess chronic cannabis use during typical consumption. Participants received Amazon gift cards of $10, $20, and $30 for completing the first, second, and third sessions, respectively.

### Procedures

The three sessions were completed within approximately one week. In the first session (45 minutes), participants completed surveys about handedness, cannabis use, alcohol use, cigarette use, anxiety, stress, mood, and cognitive function. We assessed handedness using the Edinburgh Handedness Inventory – Short Form [47,48]. To assess cannabis use, we measured: 1) frequency and quantity of use with the Daily Sessions, Frequency, Age of Onset, and Quantity of Cannabis Use Inventory (DFAQ-CU [49]), 2) cannabis dependence using the Cannabis Use Disorder Identification Test-Revised (CUDIT-R [50]), and 3) the impact of cannabis use on different aspects of daily life using the Marijuana Problem Scale (MPS [51]). We also assessed alcohol and nicotine dependence using the Alcohol Use Disorder Identification Test (AUDIT [52]) and the Fagerström Test for Nicotine Dependence (FTND [53]), respectively. To assess anxiety, mood, and stress, we used the Depression, Anxiety, and Stress Scale-21 Items (DASS-21 [54]). Lastly, we assessed self-reported cognitive impairment with the Cognitive Failures Questionnaire (CFQ [55]). We also used the Procrastination Scale [56] to assess chronic procrastination and the Need for Cognition Scale [57] to assess the tendency to enjoy effortful cognitive tasks that are reported elsewhere [58].

In the second session (90 minutes), we assessed subjective craving using the Marijuana Craving Questionnaire-Short Form (MCQ-SF [59]) following which participants performed two tasks: 1) the serial reaction time (SRT) task [60] to assess implicit motor sequence learning and 2) the Corsi block-tapping test [61] to assess visuospatial short-term and working memory. The SRT task was followed by the NASA Task Load Index (NASA-TLX [62]) to assess the amount of perceived mental and physical demand during the SRT task. Participants also completed the AX-Continuous Performance Test [63] to assess cognitive control and a transport task [64] to assess sensitivity to physical and cognitive load during decision making that are reported elsewhere [58].

In the SRT task, participants responded when a stimulus appeared in one of four squares presented in a horizontal array in the center of the computer screen. Responses were made on a QWERTY keyboard. Participants responded to each stimulus by pressing the key that spatially corresponded to the stimulus location as quickly and accurately as possible (see Fig 1A).Participants pressed the “D” key using their left middle finger for the left-most location, “F” using their left index finger for the second location, “J” using their right index finger for the third location, and “K” using their right middle finger for the right-most location. The stimulus appeared for 500 ms, and the response-to-stimulus interval was randomly selected between 500-1000 ms. The task consisted of eight blocks with 120 trials each. In the first block (B0), stimuli appeared in a random order to assess baseline reaction time (RT). In Blocks 1-4 (B1-B4), stimuli appeared in a specific sequential order consisting of 12 items that repeated 10 times in each block. In Block 5 (B5), stimuli appeared in a random order. In Block 6 (B6), stimuli appeared in the same sequence as B1-B4. This task assesses implicit motor sequence learning as participants were not informed that stimuli often appeared in the same sequence [60].

**Fig 1.**
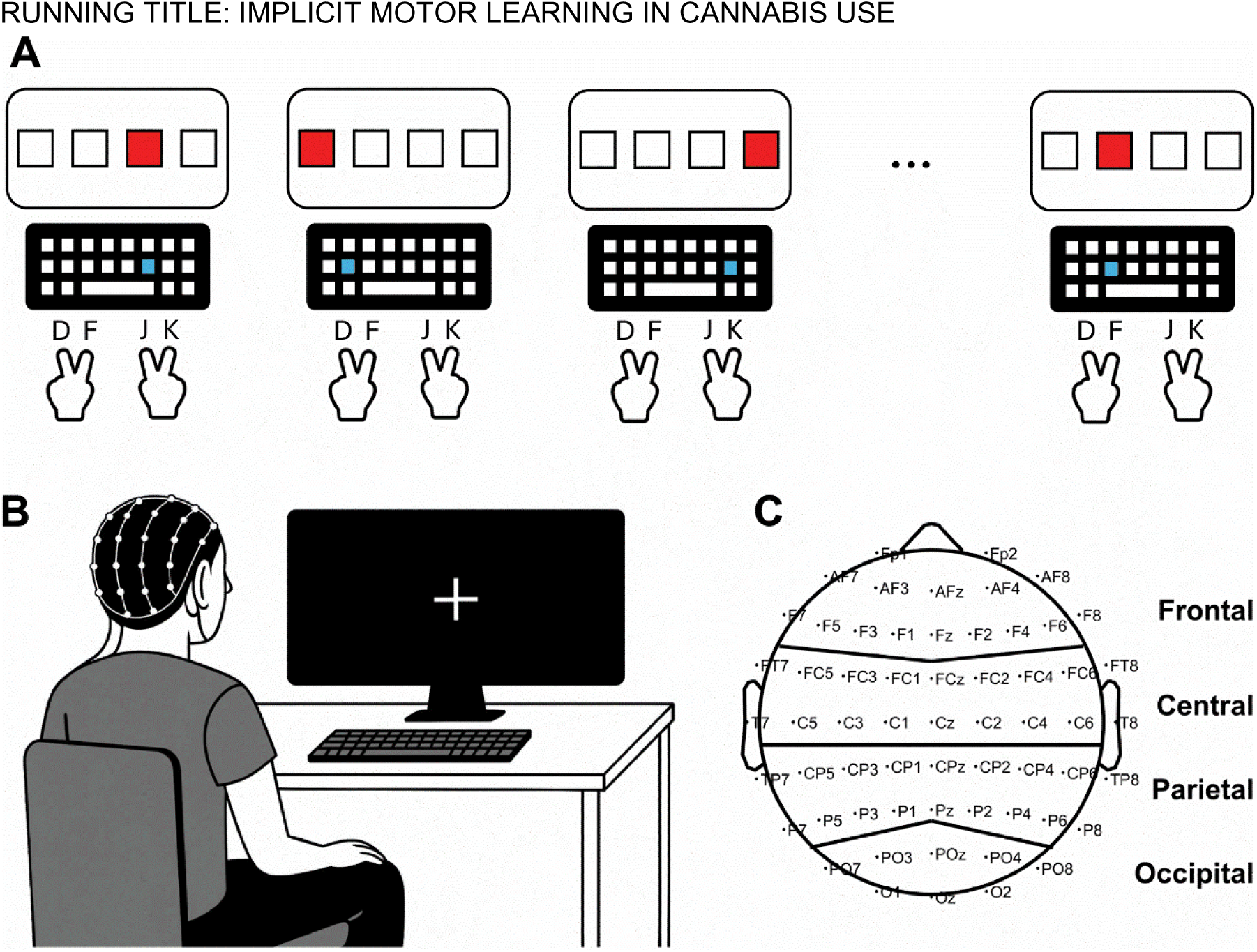
Experimental setup. A) In the serial reaction time (SRT) task, when one of the four squares on the screen turned red, the participant immediately pressed the spatially corresponding key as quickly and accurately as possible. Participants placed their fingers on the “D” (left middle finger), “F” (left index finger, “J” (right index finger), and “K” (right middle finger). B) Resting state EEG activity was recorded while participants had their eyes closed. C) The scalp montage depicts the arrangement of the 64 electrodes according to the international 10-20 system and the definition of the four cortical regions (i.e., frontal, central, parietal, and occipital regions).

In the Corsi block-tapping test, participants were presented with squares in different locations on the computer screen. In each trial, stimuli appeared in these squares in a specific order. In the forward condition, participants were instructed to click the squares in the same order. In the backward condition, participants were instructed to click the squares in the reverse order. The Corsi span score represented the largest number of items participants were able to remember in the forward condition (i.e., short-term memory) and the backward condition (i.e., working memory). The SRT task and the Corsi block-tapping test were presented using Presentation software (Neurobehavioral Systems Inc., Berkeley, CA).

Finally, in the third session (75 minutes), we assessed subjective craving again using the MCQ-SF and global cognition using the Montreal Cognitive Assessment (MoCA [65]). Afterward, we recorded eyes closed resting state EEG from each participant (see Fig 1B) with a 64-electrode ActiCap with BrainAmp amplifier (Brain Products GmbH, Gilching, Germany) with electrodes arranged according to the international 10-20 system (see Fig 1C). We used a sampling frequency of 1000 Hz and referenced to the sensor on the left earlobe. Channel impedances were below 7 kΩ. We collected three trials of two minutes each and instructed participants to close their eyes, relax, and try not to think about anything in particular.

### EEG analysis

We first processed the EEG to reduce the presence of artifacts. To reduce longitudinal drift, EEG data were high-pass filtered with a 4^th^-order zero-phase Butterworth filter at a cutoff frequency of 0.1 Hz. Brief deflections in the EEG signals, caused by sudden head movements, were reduced with artifact subspace reconstruction [66,67]. This technique removes reconstructed artifacts from time windows that contain unusually high variances in amplitude. We used a time window length of 0.5 seconds and removed artifacts from time windows that had a variance beyond 60 standard deviations compared to other data that had no deflections. Next, we removed artifacts associated with ocular, muscular, and power line activity with Independent Component Analysis (ICA) and the “ICLabel” functions [68] in the EEGLAB toolbox [69]. ICA decomposes the EEG signals into statistically independent components [70] while ICLabel automatically classifies which components are artifacts based on a large freely available database [68,71].

Then, we extracted the amount of brain wave activity as the estimated spectral power from the EEG recordings. First, we re-referenced the EEG to the common average reference. Next, we extracted the 90-second epoch that began 15 seconds after the participant was given the cue to close their eyes. Next, the linear trend from the epoch was removed to reduce spectral leakage. The Welch method was used to calculate the power spectral density (PSD), which estimates the PSD based on the average Fast Fourier Transform (FFT) solution from split segments from the epoch. The FFT was calculated in 10-second segments with a 5-second overlap, each of which was multiplied by a Hamming window using an FFT length of 8192 samples [72]. From the PSD, we extracted power from five frequency bands: delta (<4 Hz), theta (4-8 Hz), alpha (8-13 Hz), beta (13-30 Hz), and gamma (30-50 Hz). We calculated relative spectral power as a percentage of the total power (1-50 Hz) for each frequency band and grouped the EEG sensors into four regions that corresponded to major cortical areas (i.e., frontal [FP1, FP2, AF3, AFz, AF4, AF7, AF8, F1, Fz, F2, F3, F4, F5, F6, F7, and F8], central [FC1, FCz, FC2, FC3, FC4, FC5, FC6, FT7, FT8, C1, Cz, C2, C3, C4, C5, C6, T7, and T8], parietal [CP1, CPz, CP2, CP3, CP4, CP5, CP6, TP7, TP8, P1, Pz, P2, P3, P4, P5, P6, P7, and P8], and occipital [PO3, Poz, PO4, PO7, PO8, O1, Oz, and O2] regions; see Fig 1C).

### Statistical analyses

To evaluate differences in implicit motor learning, we calculated mean RTs for each block in the SRT task. We excluded incorrect responses and RTs that were greater or less than 2.5 standard deviations from each participant’s mean from the analysis [73,74]. We conducted a mixed factorial analysis of variance (ANOVA) on RT with Group (control, cannabis) as the between-subjects factor and Block (B0-6) as the within-subjects factor. If implicit learning occurred in this task, RTs should significantly decrease in the last learning block (i.e., B4) compared to the first learning block (i.e., B1) as participants learned the sequence and could respond faster by anticipating the response to the upcoming stimulus before it appeared [75,76]. Since B5 contained randomly ordered stimuli, mean RTs should significantly increase in B5 compared to B4 as participants could no longer anticipate the next stimulus. The index of motor learning [77] is quantified as an increase in RT in response to random stimuli (B5) compared to the sequenced stimuli (B4). We also expected sequenced stimuli in B6 to show significantly decreased RT compared to random stimuli in B5.

To assess differences in short-term and working memory, we performed independent *t*-tests on the forward and backward Corsi span, respectively.

To assess differences in resting state EEG, we conducted a 2 x 4 mixed-design ANOVA with the between-subjects factor of Group (control, cannabis) and the within-subject factor of Region (frontal, central, parietal, and occipital) on relative spectral power for each frequency band.

We also calculated Pearson’s correlations between variables related to cannabis use (i.e., CUDIT-R, MPS, MCQ-Session 2, MCQ-Session 3, cannabis use in the prior 30 days, years of cannabis use, age at onset of cannabis use, and lifetime cannabis use) and mood, anxiety, stress, cognition, index of motor learning from the SRT task, forward and backward Corsi span, and spectral power of cortical EEG activity in the cannabis group.

For all analyses, we used Bonferroni *post hoc* tests to account for multiple comparisons in significant effects. Any significant differences between groups on other variables such as age, education, alcohol use, or nicotine use, were included as covariates in the analyses. Statistical significance was defined at *p* < 0.05. We analyzed the data using custom scripts in MATLAB (MathWorks, Natick, MA) and SPSS (IBM, Armonk, NY).

## Results

*Demographics, cognitive-emotional assessments, and cannabis use measures* Differences in demographic, cognitive-emotional assessments, and cannabis use measures for the two groups are reported in Table 1. There were no significant differences between the groups in age [*t*(60) = 0.30, *p* = 0.77, *d* = 0.082], gender [*t*(60) =-0.44, *p* = 0.66, *d* = 0.14], education [*t*(60) = 1.5, *p* = 0.14, *d* = 0.37], or nicotine use (no participants reported nicotine use). The cannabis group reported significantly higher alcohol use [*t*(60) =-2.7, *p* = 0.010, *d* = 0.66] compared to the control group, so alcohol use was included as a covariate in subsequent analyses. For cognitive-emotional assessments, there were no differences in self-reported anxiety [*t*(60) =-1.3, *p* = 0.21, *d* = 0.31], mood [*t*(60) = 0.019, *p* = 0.99, *d* = 0.028], or stress [*t*(60) = 0.56, *p* = 0.58, *d* = 0.14]. There were also no differences in global cognition [*t*(60) =-0.61, *p* = 0.55, *d* = 0.16] or self-reported cognitive impairment [*t*(60) = 1.8, *p* = 0.081, *d* = 0.45].

The cannabis group used cannabis on significantly more days in the prior 30 days [*t*(60) =-40.8, *p* < 0.001, *d* = 8.6], for significantly more years [*t*(60) =-11.6, *p* < 0.001, *d* = 2.9], and had significantly greater lifetime cannabis use [*t*(60) =-5.2, *p* < 0.001, *d* = 1.3] compared to the control group. In addition, the cannabis group had higher MPS [*t*(60) =-5.2, *p* = 0.001, *d* = 1.3] and CUDIT-R [*t*(60) =-16.1, *p* < 0.001, *d* = 4.0] scores than the control group. Moreover, the cannabis group had significantly higher MCQ-SF scores compared to the control group [Session 2, *t*(60) =-7.0, *p* < 0.001, *d* = 1.8 and Session 3, *t*(60) =-7.8, *p* < 0.001, *d* = 2.0; the MCQ-SF was not collected in the first session].

Within the cannabis group, 10 reported using a water pipe, six reported using joints, six reported using a vaporizer, three reported using a hand pipe, another three reported using edibles, one used blunts, and one used a nectar collector. In terms of THC content, four participants reported using cannabis with an average THC content of greater than 30%, five reported using 25-30%, 12 reported using 20-24%, three reported using 15-19%, one reported using 10-14%, and five reported that they did not know.

There was a significant positive partial correlation (controlled for alcohol use) between the MPS score and anxiety (*r* = 0.30, *p* = 0.047) as well as self-reported cognitive impairment (*r* = 0.30, *p* = 0.043) in the cannabis group. In addition, age at onset of cannabis use was positively partially correlated with global cognition (*r* = 0.37, *p* = 0.013). Other correlations are included in S1 Table in the supplemental materials.

### Implicit motor learning

As we found significantly higher alcohol use in the cannabis group, we included the AUDIT score as a covariate and conducted a mixed factorial ANCOVA on RT from the SRT task. We found a main effect of Block, *F*(6,354) = 22.7, *p* < 0.001, partial *η*^2^ = 0.28, but no main effect of Group, *F*(1,59) = 0.13, *p* = 0.72, partial *η*^2^ = 0.002 or AUDIT score, *F*(1,59) = 0.23, *p* = 0.63, partial *η*^2^ = 0.004. There were also no significant interactions between Block and Group, *F*(6,354) = 0.41, *p* = 0.88, partial *η*^2^ = 0.007 or Block and AUDIT score, *F*(6,354) = 0.92, *p* = 0.48, partial *η*^2^ = 0.015. Fig 2A shows mean RTs for each block of the SRT task for both groups. *Post hoc* analyses corrected for multiple comparisons revealed a significant decrease in RT from B1 to B4 for both groups (control group, *p* = 0.001, *d* = 0.79; cannabis group, *p* < 0.001, *d* = 0.88), a significant increase in B5 compared to B4 (i.e., index of motor learning, control group, *p* < 0.001, *d* = 1.1; cannabis group, *p* = 0.008, *d* = 0.76), and a significant decrease from B5 to B6 (control group, *p* < 0.001, *d* = 1.3; cannabis group, *p* < 0.001, *d* = 0.96).

**Fig 2.**
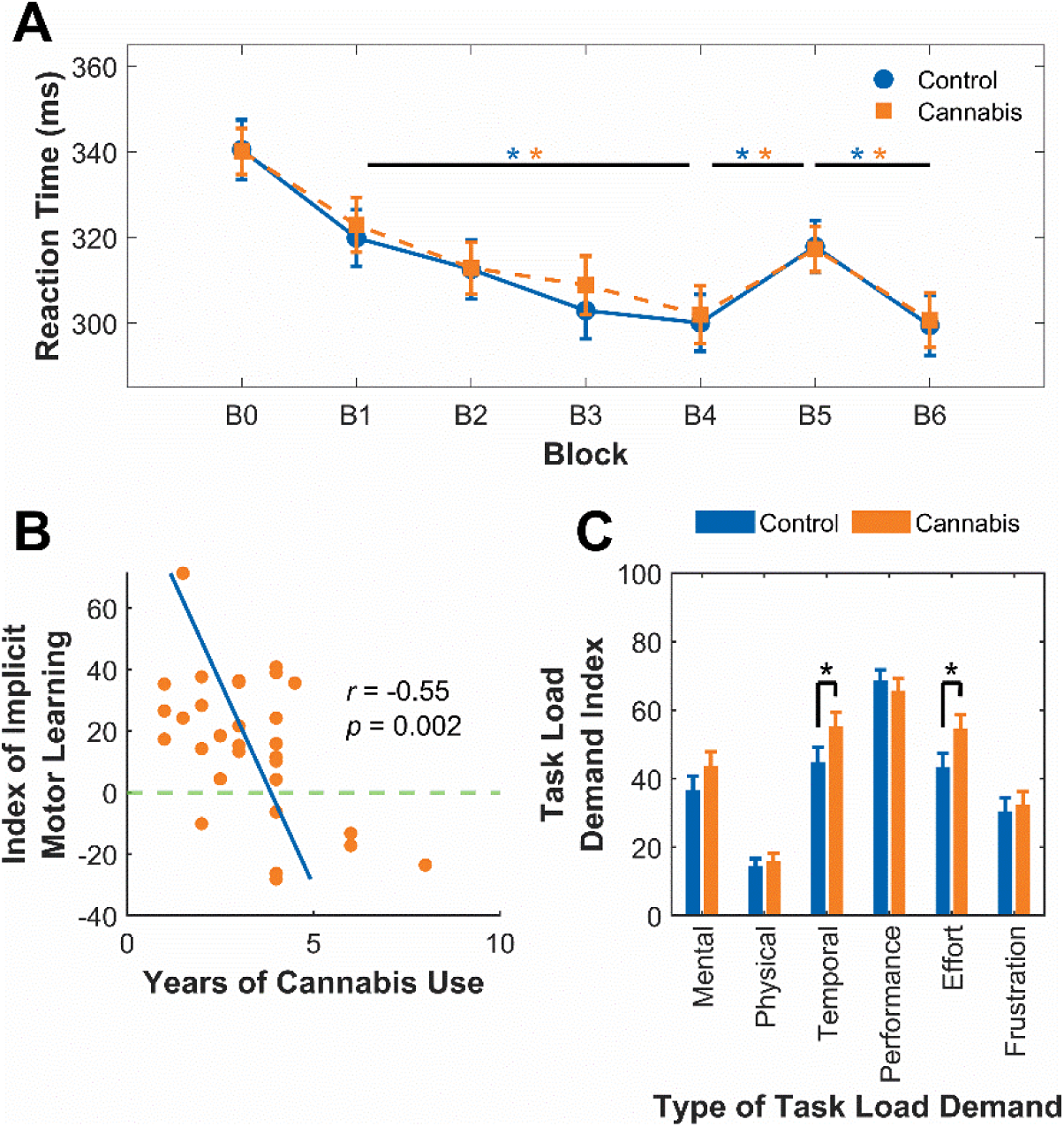
Implicit motor learning. A) Mean reaction times for correct responses in each block for the control group and the cannabis group in the serial reaction time (SRT) task. B) Individuals with more years of cannabis use showed a smaller index of implicit motor learning in the SRT task. The index of implicit motor learning in the SRT task is the difference in reaction time between Block 5 and Block 4 (i.e., a greater positive difference indicates a greater amount of learning, while a negative difference indicates no learning). The horizontal line at 0 represents a 0 ms difference between these blocks. C) The cannabis group reported feeling significantly greater temporal demand (i.e., feeling more rushed) and exerting greater effort compared to the control group when performing the SRT task. Higher task load demand index scores represent subjective reports of higher demand. The performance task load was reverse coded to match the other types of task load. Error bars indicate standard error. *Significance level of *p* < 0.05.

While there were no significant differences between the groups in the SRT task (see Fig 2A), there was a significant negative partial correlation between years of cannabis use and the index of motor learning (*r* =-0.55, *p* = 0.002) in the cannabis group, shown in Fig 2B.

To evaluate perceived mental and physical demand of the SRT task, we compared ratings on the NASA-Task Load Index, shown in Fig 2C. The cannabis group reported exerting significantly greater effort [*t*(60) =-2.0, *p* = 0.025, *d* = 4.8] and feeling more rushed [*t*(60) =-1.7, *p* = 0.046, *d* = 5.0] during the task compared to the control group. There were no significant differences in perceived mental demand [*t*(60) =-1.2, *p* = 0.12, *d* = 4.9], physical demand [*t*(60) =-0.42, *p* = 0.34, *d* = 2.5], successful performance [*t*(60) =-0.64, *p* = 0.26, *d* = 3.9], or frustration [*t*(60) =-0.34, *p* = 0.37, *d* = 4.5] between groups.

### Resting state EEG

In the delta, theta, alpha, and beta bands, there was a significant main effect of Region [delta, *F*(3,177) = 103.0, *p* < 0.001, partial *η*^2^ = 0.64; theta, *F*(3,177) = 14.3, *p* < 0.001, partial *η*^2^ = 0.20; alpha, *F*(3,177) = 81.8, *p* < 0.001, partial *η*^2^ = 0.58; beta, *F*(3,177) = 12.8, *p* < 0.001, partial *η*^2^ = 0.18], but no significant main effect of Group and no significant interactions. Relative spectral power scalp maps for each frequency band for both groups are depicted in Fig 3. In the gamma band, there were no significant main effects, but there was a significant interaction between Region x Group, *F*(3,177) = 4.2, *p* = 0.007, partial *η*^2^ = 0.066. *Post hoc* analyses indicated no significant differences between regions in the control group, but in the cannabis group, there was significantly higher gamma power in the central region compared to the frontal (*p* < 0.001, *d* = 1.3), parietal (*p* < 0.001, *d* = 1.1), and occipital (*p* = 0.004, *d* = 1.2) regions.

**Fig 3.**
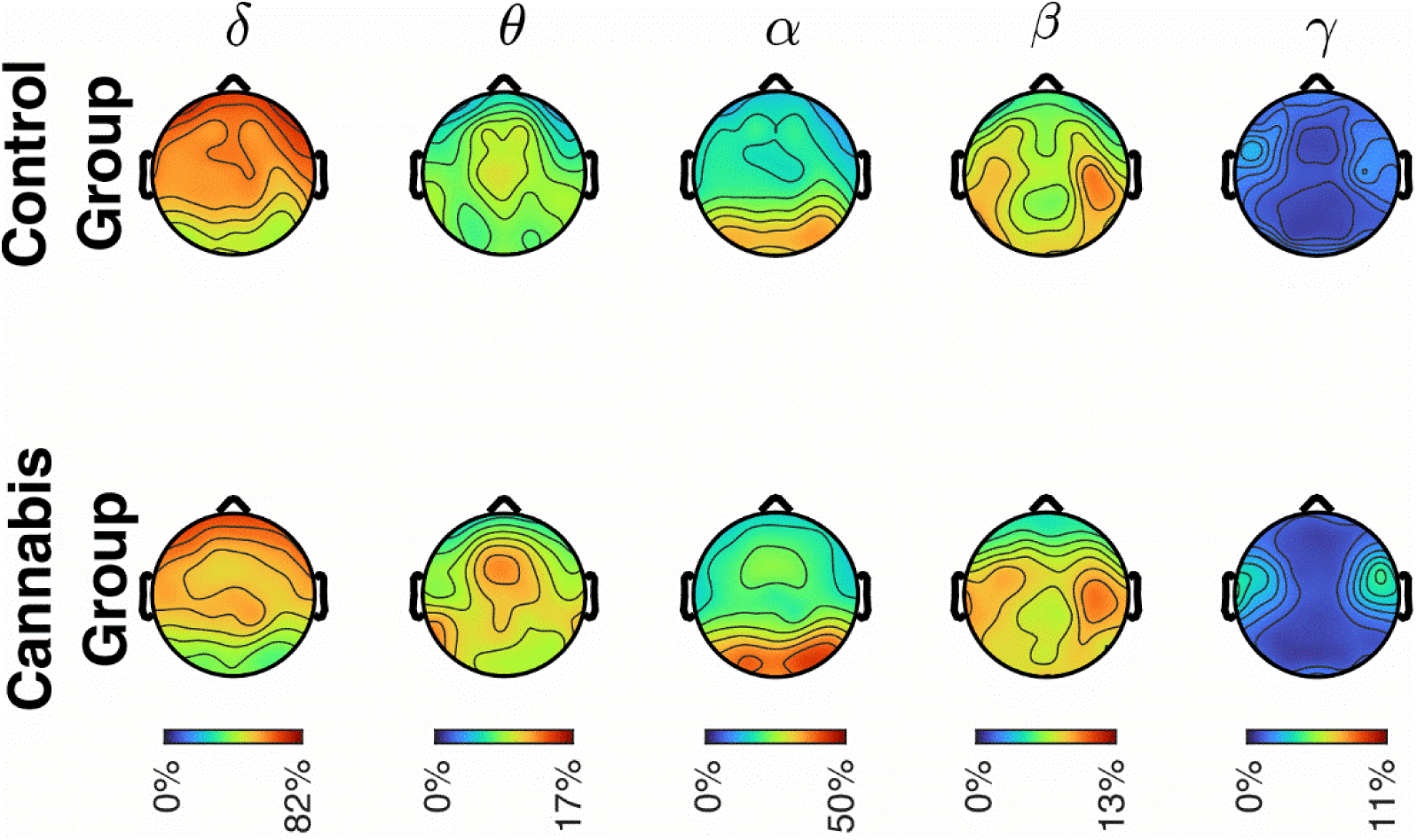
Scalp map depicting relative spectral power for each frequency band for the control and cannabis groups. There were no significant differences between the groups for any frequency band. Colors represent high (red) or low (blue) spectral power.

We predicted that the cannabis group would exhibit supranormal EEG spectral power (i.e., greater spectral power in the beta and gamma bands and less spectral power in the delta, theta, and alpha bands), but we did not find group level differences. However, within the cannabis group, we found significant correlations between cannabis use measures and beta, gamma, and delta frequencies (see Fig 4). Specifically, age at onset of cannabis use was negatively partially correlated with beta activity in the central (*r* = - 0.43, *p* = 0.014) regions. In addition, cannabis use in the prior 30 days was positively partially correlated with gamma activity in the frontal (*r* = 0.62, *p* < 0.001), central (*r* = 0.43, *p* = 0.019), and occipital (*r* = 0.48, *p* = 0.008) regions. Finally, subjective craving assessed during the third session was negatively partially correlated with delta activity in the parietal (*r* =-0.37, *p* = 0.049) and occipital (*r* =-0.38, *p* = 0.041) regions, positively partially correlated with beta activity in the frontal region (*r* = 0.48, *p* = 0.007), and positively partially correlated with gamma activity in the frontal (*r* = 0.45, *p* = 0.014) and occipital (*r* = 0.51, *p* = 0.005) regions. A subset of the above correlations are depicted in Fig 4 and the remaining are in the supplementary materials (S2 Fig).

**Fig 4.**
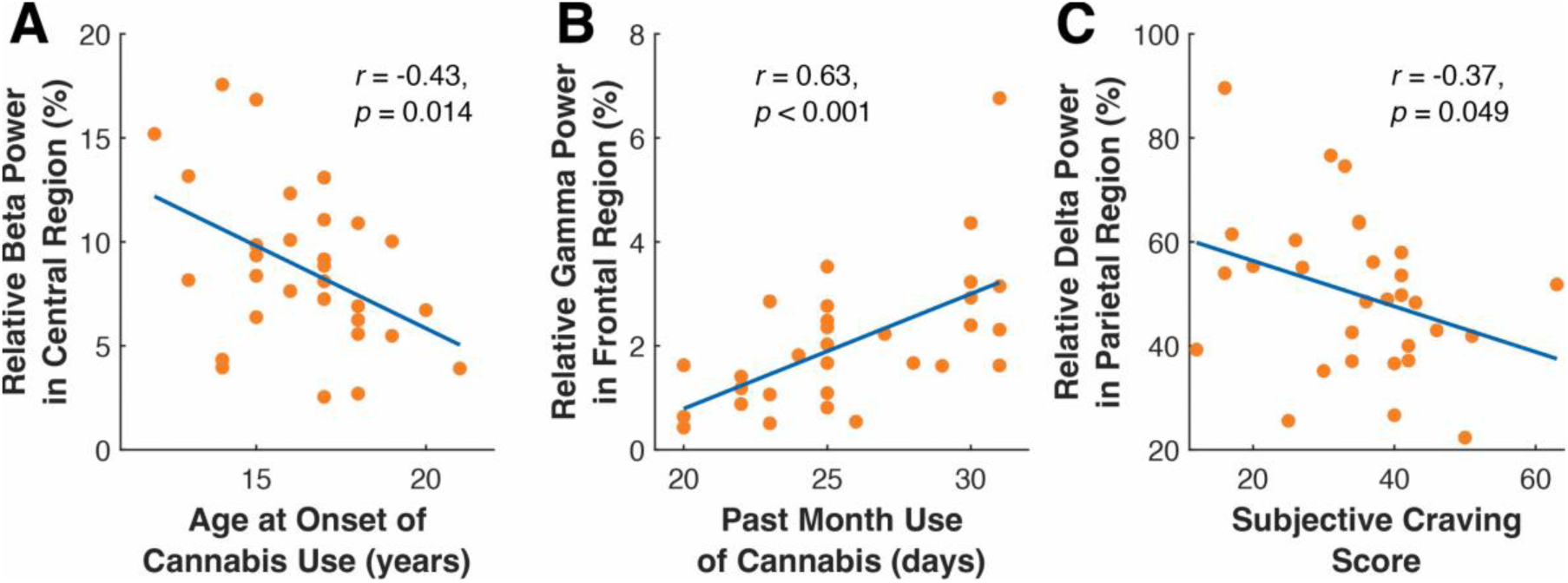
Correlation between cannabis use measures and spectral power of cortical EEG activity. A) Higher beta activity in the central region was associated with an earlier age of onset of cannabis use. B) Higher gamma activity in the frontal (shown), central (not shown), and occipital (not shown) regions was associated with higher cannabis use in the prior 30 days. C) Lower delta activity in the parietal (shown) and occipital (not shown) regions was associated with higher subjective craving of cannabis (measured via the Marijuana Craving Questionnaire-Short Form; MCQ-SF).

### Corsi block-tapping test

The cannabis group had a significantly shorter forward (i.e., visuospatial short-term memory, *t*(60) = 2.5, *p* = 0.017, *d* = 1.2) and backward (i.e., visuospatial working memory, *t*(60) = 2.9, *p* = 0.006, *d* = 1.5) span compared to the control group (see Fig 5). Additionally, subjective craving assessed in the second session was negatively partially correlated with the backward Corsi span (*r* =-0.37, *p* = 0.049) in the cannabis group.

**Fig 5.**
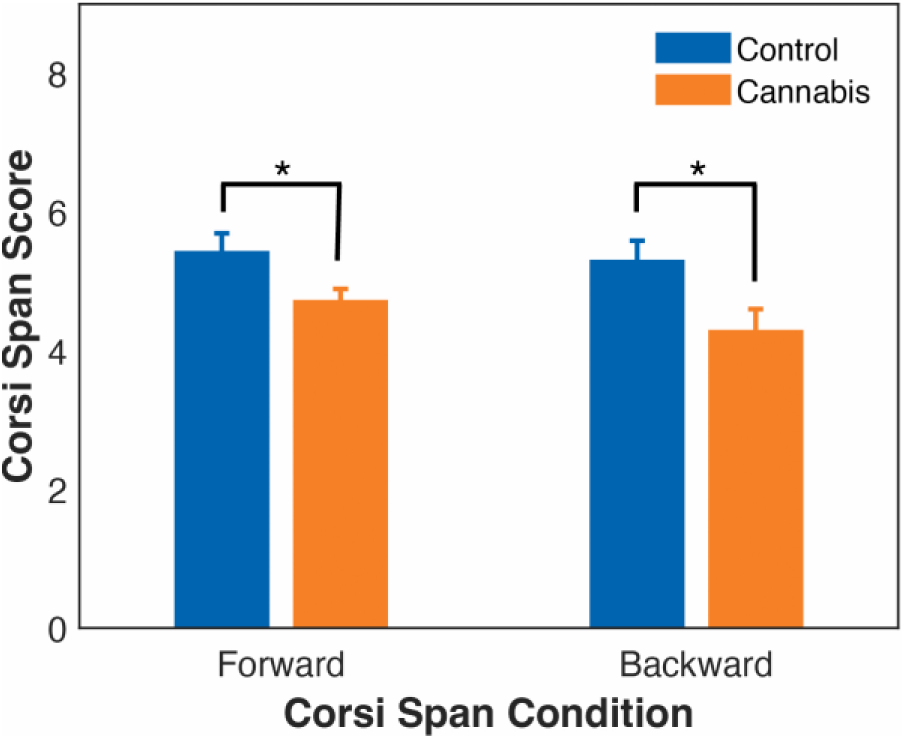
Memory span in the control and cannabis groups measured by the Corsi block-tapping test. The cannabis group had significantly lower Corsi span scores in both the forward (i.e., visuospatial short-term memory) and backward (i.e., visuospatial working memory) conditions. *Significance level of *p* < 0.05.

## Discussion

We examined the effect of chronic cannabis use on implicit motor learning, resting state cortical EEG activity, and visuospatial short-term and working memory. Our results indicate that more years of cannabis use was associated with a smaller index of implicit motor learning, younger age at onset of use was associated with increased beta oscillations, increased past month use was associated with increased gamma oscillations, and higher subjective craving was associated with decreased delta oscillations. In addition, we found significantly reduced visuospatial short-term and working memory spans in the cannabis group. Finally, the cannabis group reported exerting greater effort during the implicit motor learning task, which may have contributed to their performance being comparable to the control group and greater effort may have been necessary to compensate for their increased resting state cortical EEG activity. These findings suggest that supranormal resting state EEG activity may increase cortical noise, interfere with visuospatial short-term and working memory, and affect implicit motor learning such that greater effort is required for motor performance to be comparable to the control group.

### Implicit motor learning was impaired in individuals with longer chronic cannabis use

We found intact implicit motor learning in the cannabis group which was no different than the control group. However, the cannabis group reported that they exerted significantly greater effort and felt more rushed while performing the SRT task. Feeling more rushed may be a consequence of the greater effort exerted during the task. This greater effort may have been necessary for the cannabis group to perform at a level that was comparable to that of the control group. While the amotivation syndrome hypothesis [78,79] suggests that acute cannabis use causes apathy [80,81], recent studies found that individuals who use cannabis chronically may be more willing to exert more effort to improve performance [58,81–87]. Although we did not find group-level differences in implicit motor learning, there were individual differences within the cannabis group.

Specifically, a smaller index of implicit motor learning was associated with longer chronic cannabis use, indicating a potential link to reduced implicit motor learning in these individuals. This finding, together with the increased resting state beta activity that was associated with an earlier onset of cannabis use, suggests that longer chronic cannabis use may impact the corticostriatal pathway that plays a critical role in motor learning [88,89]. These results complement recent evidence demonstrating that chronic cannabis use impairs visuomotor adaptation (i.e., successful application of previously well-learned motor skills to new contexts) [90,91]. However, motor adaptation relies on the corticocerebellar circuit, whereas implicit motor sequence learning relies on the corticostriatal circuit [88]. Our novel finding suggests that the corticostriatal pathway may also be impaired by longer chronic cannabis use, advocating a more widespread impact of chronic cannabis use on the motor system. To fully understand how cannabis affects implicit motor learning, future studies may systematically explore the distinct effects of acute versus chronic use. Importantly, implicit motor learning is essential for acquiring and automating motor skills that we perform in our daily lives, such as using new tools and technologies, navigating unfamiliar environments, and adapting to changing task demands. When these processes are impaired, individuals may struggle to learn new motor skills or adapt to novel situations, particularly when stressed, distracted, or fatigued. This, in turn, can restrict performance, compromise daily functioning, threaten safety, and jeopardize independence.

### Supranormal resting state EEG may underlie implicit motor learning impairment

We found that within the cannabis group, cannabis use measures were correlated with supranormal resting state neural oscillations. Specifically, a younger age of onset of cannabis use was associated with higher beta activity, and higher use of cannabis in the prior 30 days was associated with higher gamma activity. The shift in cortical activity may be due to the interaction between THC and CB1 receptors that can modulate the balance between excitatory and inhibitory postsynaptic activity and regulate local cortical excitability [92]. Specific to gamma activity, THC exposure can decrease the release of GABA in the prefrontal cortex and disrupt cortical gamma activity in adolescents [93].

Furthermore, another study reported that individuals who use cannabis chronically exhibited increased resting state beta activity [41], which is also consistent with our results. Tempel and colleagues found that when retrieving motor sequences from memory, increased beta power predicted motor forgetting [94]. Importantly, we found increased beta activity in the central areas, which are critical for motor control [95–97]. While our results only reflect changes in resting state, the effect cannabis use has on resting beta activity should be considered in future studies that examine task-related activity.

Our finding of supranormal cortical EEG activity during resting state may reflect increased cortical noise in the resting functional organization of the brain. Earlier studies have found that increased cortical noise during the baseline period before stimulus presentation in an oddball task was associated with acute THC use in individuals who abstained from cannabis for an average of 445.7 ± 846.6 days [98] and in individuals who abstained from cannabis 12 hours prior to the session [99]. Thus, these earlier studies have demonstrated that acute THC exposure after abstinence alters cortical activity. Our findings extend this evidence by suggesting that increased cortical noise may also occur with chronic cannabis use without abstinence. It is unclear whether these effects persist with abstinence from cannabis or if they endure long-term. Future studies may consider a longitudinal approach to understand the directionality of these effects and parse whether short-term or long-term abstinence allows for recovery of motor learning.

### Lower visuospatial short-term and working memory capacity in the cannabis group

A critical aspect of motor learning is the ordering of action sequences; this acquisition requires short-term and working memory to combine individual actions into complex motor behavior [8]. Visuospatial working memory, in particular, is critical for optimum motor control [100] and is positively correlated with the rate of motor learning [101]. We found that individuals in the cannabis group had significantly reduced forward and backward Corsi spans compared to those in the control group, suggesting that cannabis use was associated with reduced visuospatial short-term and working memory. This finding is consistent with prior studies reporting that more frequent cannabis use was associated with poorer working memory and reduced hippocampal volume [102].

Furthermore, animal studies have directly demonstrated the impact of THC on spatial working memory. Rats given THC exhibited dose-dependent impaired short-term and working memory in a water maze task [103,104]. In addition, D1 receptors in the prefrontal cortex are involved in working memory and are stimulated by cannabis [15,105]. When D1 receptors were activated at a higher level than is typical, spatial working memory was impaired in rhesus monkeys [106] and rodents [107]. This evidence is consistent with our findings that there was significantly reduced visuospatial short-term and working memory in the cannabis group. In addition, the reduced visuospatial memory capacity in the cannabis group may have contributed to their need for greater effort during the SRT task compared to the control group.

### Limitations and future directions

Our results must be interpreted within the limitations of the current study. Our primary conclusions are based on correlations between cannabis use measures, index of motor learning, and resting state cortical EEG activity. While these provide a basis for the effect of cannabis, further research is needed to draw causal conclusions. For example, future studies may assess whether there are group differences between individuals who used cannabis for a greater number of years compared to those who used cannabis for fewer years. We also asked participants to use cannabis as they normally would to capture their typical daily functioning, rather than during an imposed abstinence period. While this approach preserved ecological validity, it also confounded the effects of acute and chronic use. However, including an abstinence period may increase the level of craving and withdrawal which may impact implicit motor learning and resting state cortical activity. Furthermore, previous studies have reported that anxiety tends to increase during abstinence from cannabis, particularly within the first 24 hours [108–113]. Since heightened anxiety can impair cognitive function [114–116], it may confound interpretations of the effects of chronic cannabis use. Nevertheless, it is important for future studies to explore the differences between acute and chronic effects on implicit motor learning to better understand the nuanced impact of cannabis. In addition, we did not control the amount of cannabis use beyond the inclusion criterion of at least four uses per week for at least one year. Consequently, variability in consumption patterns may have influenced the findings. Future research may examine the impact of dosage and product variability (e.g., THC concentrations, forms of consumption) to better understand their impact on implicit motor learning and resting state EEG activity.

Our EEG analysis focused on relative spectral power to directly measure neural oscillatory activity and to assess differences in the distribution of this oscillatory activity across different frequencies. Future studies could utilize source localization techniques (e.g., standardized low resolution brain electromagnetic tomography (sLORETA [117], dipole source localization [118], and beamforming [119]) to help identify cortical origins of EEG signals and provide insight into how specific cortical areas may be affected by cannabis use. In addition, techniques such as functional connectivity could be used to assess interhemispheric communication and global brain connectivity [120] which may elucidate how cannabis use alters cortico-cortical communication between regions, particularly those involved in motor control and working memory, including the prefrontal cortex, primary motor cortex, premotor cortex, supplementary motor area, and parietal regions.

Finally, both groups in our sample contained a large proportion of female participants. As prior studies have indicated sex-related differences in neural and behavioral responses to cannabis [121–123], future studies may explore whether these differences impact implicit motor learning and resting state cortical activity.

## Conclusions

We found that longer chronic cannabis use was associated with impaired implicit motor learning. Furthermore, the cannabis group exhibited significantly lower visuospatial short-term and working memory compared to the control group. Such functional impairments may be associated with altered resting state neural oscillations we observed in this study. We found that an earlier age of onset of use was associated with increased resting state beta activity, greater use in the prior month was associated with increased resting state gamma activity, and a higher subjective craving was associated with decreased resting state delta activity. We suggest that the increase in resting state neural oscillations may reflect increased cortical noise that may disrupt cognitive and motor processing. Collectively, these findings suggest that longer chronic cannabis use may affect the corticostriatal network, which plays an important role in implicit motor learning, revealing a more widespread impact of chronic cannabis use on the motor system. Our findings highlight the complex nature of the effect of chronic cannabis use on cortical activity, cognition, and implicit motor learning with important considerations for understanding the full impact of cannabis on brain health, instrumental activities of daily living, and the consequent impact on public health.

## Author Contributions

SP and LRF designed the study. SP and AYP analyzed the data. SP lead the writing of the manuscript. All authors collected data and reviewed/contributed to ideas contained in the manuscript.

## Supporting information

**S1 Table.** Partial correlations (controlled for alcohol use) between variables related to cannabis use and cognitive-emotional assessments, the serial reaction time (SRT) task, the Corsi block-tapping task, and spectral power in the cannabis group.

**S2 Fig. Correlation between cannabis use measures and spectral power of cortical EEG activity (in additional regions not shown in figure in manuscript).** Higher gamma activity in the A) central and B) occipital regions was associated with higher cannabis use in the prior 30 days. C) Lower delta activity in the occipital region was associated with higher subjective craving of cannabis (measured via the Marijuana Craving Questionnaire-Short Form; MCQ-SF). D) Higher beta activity in the frontal region was also associated with higher subjective craving.

Higher gamma activity in the E) frontal and F) occipital regions was associated with higher subjective craving as well.

