## Supplemental Information for "Longer chronic cannabis use in humans is associated with impaired implicit motor learning and supranormal resting state cortical activity"

### Supporting Information

**Table S1.** Partial correlations (controlled for alcohol use) between variables related to cannabis use and cognitive-emotional assessments, the serial reaction time (SRT) task, the Corsi block-tapping task, and spectral power in the cannabis group.

| | Lifetime cannabis use | Years of cannabis use | Past month cannabis use | Age at onset of cannabis use | CUDIT-R | MPS | MCQ-SF (Session 2) | MCQ-SF (Session 3) |
|---|---|---|---|---|---|---|---|---|
| **Cognitive-Emotional Assessments** | | | | | | | | |
| DASS-21 | | | | | | | | |
| Depression | 0.15 | -0.017 | 0.075 | -0.13 | 0.20 | 0.26 | - | - |
| Anxiety | 0.13 | 0.14 | 0.14 | -0.27 | 0.15 | 0.30* | - | - |
| Stress | 0.14 | -0.21 | -0.11 | -0.059 | -0.084 | 0.26 | - | - |
| CFQ | -0.022 | -0.23 | -0.16 | -0.20 | -0.003 | 0.30* | - | - |
| MoCA | -0.17 | -0.086 | 0.045 | 0.37* | -0.011 | -0.067 | - | - |
| **Serial Reaction Time Task** | | | | | | | | |
| Index of motor learning | -0.081 | -0.55* | -0.15 | 0.33 | -0.022 | -0.13 | -0.15 | - |
| **Corsi Block-Tapping Task** | | | | | | | | |
| Forward span | -0.14 | -0.19 | 0.013 | 0.070 | -0.13 | -0.16 | -0.18 | - |
| Backward span | -0.093 | -0.046 | -0.021 | 0.12 | -0.092 | -0.15 | -0.37* | - |
| **EEG** | | | | | | | | |
| Delta | | | | | | | | |
| Frontal | 0.17 | 0.15 | -0.020 | 0.14 | 0.084 | 0.082 | - | -0.32 |
| Central | 0.074 | 0.14 | -0.18 | 0.27 | 0.038 | 0.032 | - | -0.29 |
| Parietal | -0.012 | 0.13 | -0.16 | 0.19 | -0.002 | 0.010 | - | -0.37* |
| Occipital | -0.065 | 0.033 | -0.23 | 0.19 | -0.21 | -0.23 | - | -0.38* |
| Theta | | | | | | | | |
| Frontal | -0.21 | -0.18 | -0.040 | -0.021 | -0.36 | -0.19 | - | 0.21 |
| Central | -0.16 | -0.15 | -0.017 | -0.070 | -0.35 | -0.18 | - | 0.19 |
| Parietal | -0.17 | -0.19 | -0.049 | -0.069 | -0.30 | -0.11 | - | 0.21 |
| Occipital | -0.19 | -0.23 | -0.059 | 0.009 | -0.33 | -0.094 | - | 0.22 |
| Alpha | | | | | | | | |
| Frontal | -0.17 | -0.069 | -0.079 | -0.13 | 0.018 | -0.061 | - | 0.17 |
| Central | -0.068 | -0.040 | 0.054 | -0.13 | -0.018 | -0.092 | - | 0.16 |

| | | | | | | | | |
|---|---|---|---|---|---|---|---|---|
| Parietal | 0.061 | -0.053 | 0.160 | -0.092 | 0.096 | -0.024 | - | 0.27 |
| Occipital | 0.085 | 0.077 | 0.16 | -0.15 | 0.30 | 0.21 | - | 0.19 |
| Beta | | | | | | | | |
| Frontal | 0.062 | -0.12 | 0.27 | -0.38 | 0.11 | 0.081 | - | 0.49* |
| Central | 0.12 | -0.10 | 0.26 | -0.44* | 0.23 | 0.24 | - | 0.34 |
| Parietal | 0.15 | -0.021 | 0.20 | -0.37 | 0.20 | 0.22 | - | 0.30 |
| Occipital | 0.21 | -0.042 | 0.26 | -0.22 | 0.22 | 0.26 | - | 0.33 |
| Gamma | | | | | | | | |
| Frontal | 0.19 | -0.11 | 0.62* | -0.20 | 0.20 | 0.21 | - | 0.45* |
| Central | 0.015 | -0.14 | 0.43* | -0.24 | 0.27 | 0.16 | - | 0.16 |
| Parietal | 0.12 | 0.074 | 0.26 | -0.24 | 0.30 | 0.19 | - | 0.018 |
| Occipital | 0.17 | -0.076 | 0.48* | -0.19 | 0.057 | 0.11 | - | 0.51* |

Abbreviations: CUDIT-R, Cannabis Use Disorder Identification Test-Revised; MPS, Marijuana Problem Scale; MCQ-SF, Marijuana Craving Questionnaire-Short Form, DASS-21, Depression, Anxiety, and Stress Scale – 21 Items; CFQ, Cognitive Failures Questionnaire; MoCA, Montreal Cognitive Assessment.
*Indicates $p < 0.05$

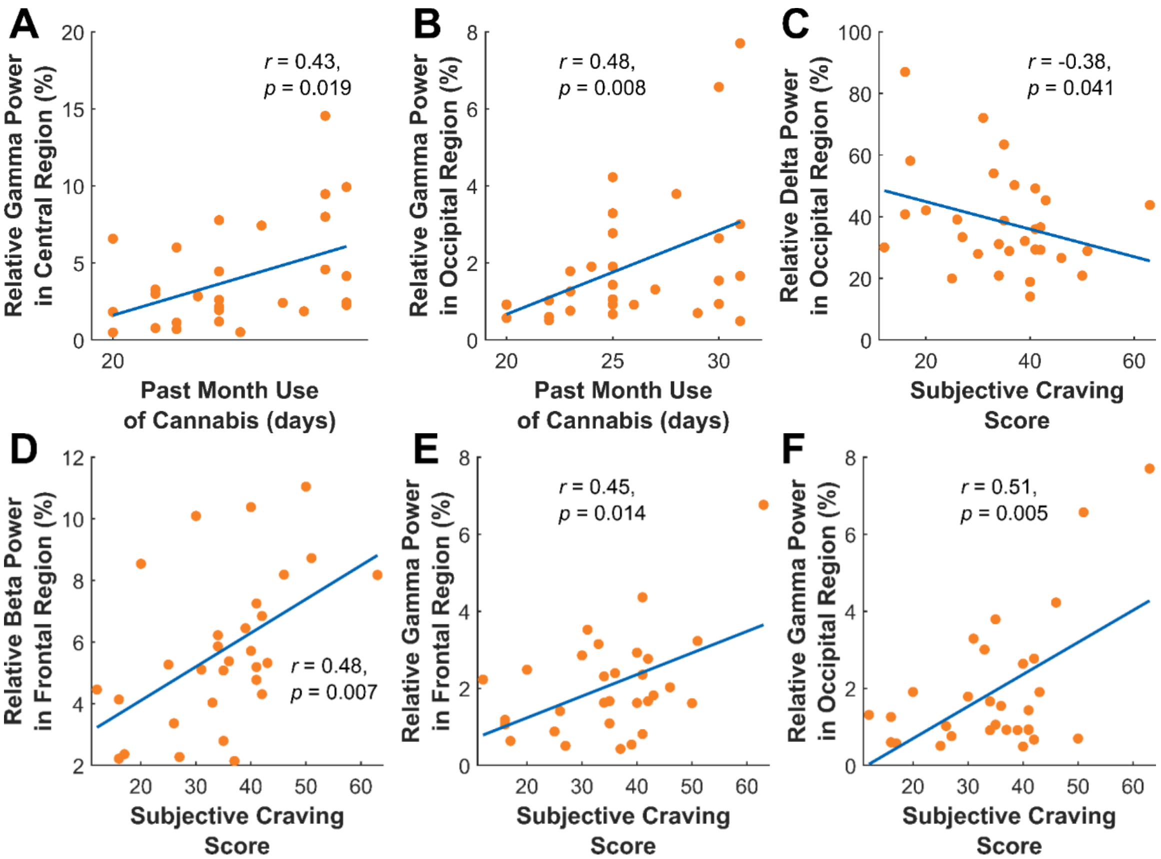


**Fig S2: Correlation between cannabis use measures and spectral power of cortical EEG activity (in additional regions not shown in figure in manuscript).** Higher gamma activity in the A) central and B) occipital regions was associated with higher cannabis use in the prior 30 days. C) Lower delta activity in the occipital region was associated with higher subjective craving of cannabis (measured via the Marijuana Craving Questionnaire-Short Form; MCQ-SF). D) Higher beta activity in the frontal region was also associated with higher subjective craving. Higher gamma activity in the E) frontal and F) occipital regions was associated with higher subjective craving as well.
